# TR-107 and Capivasertib Synergistically Inhibit Pancreatic Cancer Organoid Growth In Vitro

**DOI:** 10.64898/2026.11.13.766013

**Authors:** Joseph F. LaComb, Aisharja Mojumdar, Catalina Vera, Sahaana Kesavan, Mia Kotikovski, Lee M. Graves, Edwin J. Iwanowicz, Agnieszka B. Bialkowska

## Abstract

Pancreatic ductal adenocarcinoma (PDAC) is a leading cause of cancer-related mortality in the United States, highlighting the need for effective therapeutic strategies. Oxidative phosphorylation (OXPHOS) has emerged as a promising therapeutic target due to metabolic reprogramming and the pro-survival role of mitochondria in PDAC. The mitochondrial ClpXP complex, composed of the AAA+ ATPase ClpX and the peptidase ClpP, regulates mitochondrial homeostasis through protein degradation. Madera Therapeutics has developed novel ClpP activators, including TR-107. Here, we demonstrate that TR-107 reduces viability in four PDAC organoids and four cell lines at low-nanomolar concentrations while decreasing ClpX expression in a dose-dependent manner. Transcriptomic analysis revealed alterations in ferroptosis, amino acid metabolism, and HIF-1α signaling pathways. Notably, TR-107 retained its efficacy and mechanism of action under hypoxic conditions. Furthermore, TR-107 synergized with the AKT inhibitor capivasertib to suppress cell viability and modulate downstream signaling pathways. Collectively, these findings identify TR-107 as a promising therapeutic candidate that targets the mitochondrial ClpXP complex, disrupts mitochondrial function, and induces cellular stress and metabolic dysregulation. These effects may provide opportunities for combination therapies to improve treatment outcomes in PDAC.

## Introduction

Pancreatic cancer is the fourth deadliest cancer in the United States and is projected to become the second cause of cancer-related deaths by 2023 [1–3]. The majority of patients are diagnosed with locally advanced or metastatic disease due to a lack of early screening and delayed onset of symptoms [4–6]. These patients are not candidates for surgical resection [7]. The conventional treatment options, while not very effective, include Gemcitabine/nab-Paclitaxel [8] or FOLFIRINOX (5-Fluorouracil, Leucovorin, Irinotecan, Oxaliplatin) chemotherapy [9, 10]. Late-stage diagnosis, combined with ineffective treatments and chemoresistance [6, 11, 12], leads to poor prognosis with a median five-year survival of less than eight percent [13, 14]. Moreover, surgical resection adds limited benefit, increasing the overall median survival from 6.7-11.1 months in non-resectable patients [8, 9] to 25-28 months in patients receiving combination surgery and chemotherapy [15]. Thus, novel therapies are urgently needed to address this dismal outcome.

Metabolic reprogramming promotes cancer cell survival and proliferation in many aggressive cancers. In pancreatic ductal adenocarcinoma (PDAC), this plasticity is predominantly driven by oncogenic mutations and the tumor microenvironment (TME) [16–19]. *KRAS* mutations, which occur in more than 90% of PDAC cases, increase glycolysis, amino acid metabolism, and macropinocytosis of the extracellular milieu by activating downstream signaling pathways [17, 18]. Subsequently, the TME contributes to metabolic remodeling and chemoresistance through angiogenesis and fibrosis. PDAC cells may rely on both glycolysis and oxidative phosphorylation (OXPHOS) for energy production and biosynthesis in such conditions [20, 21]. A hallmark of PDAC, dense stroma and decreased vasculature create a hypoxic, nutrient-poor gradient. PDAC cells rely less on OXPHOS for energy production by upregulating glycolysis and proline metabolism in these conditions [19]. Disruption of metabolism, including OXPHOS, sensitizes cancers to chemotherapies while impairing tumorigenesis [22–25]. Thus, mitochondria have become an intriguing target for novel therapeutics.

The ClpXP complex is a protease that degrades damaged and misfolded proteins within the mitochondrial matrix. It comprises a caseinolytic mitochondrial matrix peptidase proteolytic subunit (ClpP) and a caseinolytic mitochondrial matrix peptidase chaperone subunit X (ClpX). ClpP is a conserved, tetradecameric mitochondrial serine protease, whereas ClpX is a hexameric AAA+ (<u>A</u>TPases <u>a</u>ssociated with diverse cellular <u>a</u>ctivities) ATPase chaperone [26]. The ATP-dependent activity of ClpX allows for recognition, unfolding, and translocation of damaged or misfolded proteins to the ClpP proteolytic chamber for degradation [26, 27]. Specifically, subunits of the TCA cycle, OXPHOS, amino acid metabolism, and mitochondrial transcription and translation factors are known ClpP substrates [28]. Dysregulated ClpP activity perturbs mitochondrial function, including OXPHOS, indicating that ClpXP is essential in maintaining mitochondrial homeostasis [29–32]. ClpX and ClpP expression varies between cancers. In PDAC, high ClpP expression is positively correlated with suppressed tumorigenesis and longer survival [22, 32]. ClpX, while not as well characterized, is negatively correlated with favorable prognosis [33] (Human Protein Atlas https://www.proteinatlas.org/ENSG00000166855-CLPX/pathology/pancreatic+cancer). The ClpXP complex may therefore be a mechanism by which novel therapies act on mitochondria and cellular respiration.

Acyldepsipeptides (ADEPs) were the first discovered agonists of ClpP in multidrug-resistant bacteria [34]. Crystallography studies revealed that ADEPs bound a hydrophobic pocket in the N-terminus of ClpP [35]. This interaction displaces ClpX and induces unregulated ClpP activity [35]. As part of the National Cancer Institute’s Diversity Set II screen, the imipridone ONC201 was identified as a potential inducer of TRAIL transcription [36]. ONC201 was found to have tumor suppressor effects in multiple cancers [37]. Further investigation showed that it disrupted mitochondrial function and OXPHOS by directly activating ClpP [38–40]. ONC201 ultimately advanced to more than 25 clinical trials for PDAC, colorectal cancer (CRC), glioblastoma, and Non-Hodgkin Lymphoma, among others, despite an unclear mechanism of action. Subsequently, Madera Therapeutics designed and synthesized a family of ONC201 analogs termed TR compounds. Several TR analogs are 50-100 times more effective against cancer cell proliferation. They have enhanced ClpP activation; exhibiting a 10-100-fold increase in ClpP activity and up to a 10-fold greater ClpP binding affinity compared to the parent ONC201 [38]. More specifically, TR-107 inhibited mitochondrial protein expression and OXPHOS via ClpP activation at low-nanomolar concentrations in CRC [41] and triple-negative breast cancer (TNBC) cells [42]. The safety and efficacy (including tumor inhibition, bioavailability, and half-life) of these analogs compared to ONC201 have been reported in mouse xenograft models [42].

We now demonstrate that TR-107 has low nanomolar IC_50_ concentrations in four patient-derived PDAC organoids and four well-established cell lines. TR-107 antagonizes ClpX via unregulated ClpP proteolytic activity in a dose-dependent manner. Transcriptomic analysis revealed aberrant gene expression after 48 hours of treatment, including increased amino acid metabolism, dysregulated ferroptosis, and suppression of HIF-1α signaling. Finally, we show that TR-107 synergized with the AKT inhibitor Capivasertib against PDAC viability while affecting ClpX and AKT expression, respectively.

## Results

### TR-107 inhibits PDAC viability at low nanomolar concentrations

The inhibitory efficacy of TR-107 was assessed in pancreatic cancer models. Human PDAC organoids hF31, hF37, hF44, and hT1 were seeded in Matrigel and treated with log-fold concentrations of the TR-107 or DMSO (vehicle control) for 72 hours. A marked decrease in viability was observed between 100 nM and 1 µM for hF31, and between 10 nM and 100 nM for hF37, hF44, and hT1 as qualitatively assessed by microscopy (Fig. 1). Viability of the organoids and human PDAC cell lines (BxPC-3, Capan-2, MIA PaCa-2, and PANC-1) was subsequently measured using the CellTiter-Glo luciferase assay. IC_50_ values were calculated using log(inhibitor) vs response – variable slope (four parameters) nonlinear fit. In Capan-2, MIA PaCa-2, and PANC-1, the IC_50_ values for TR-107 were 14 nM, 75 nM, and 1 nM, respectively (Sup. Fig. 1a). For BxPC-3, the IC_50_ was ambiguous due to the nonlinear fit (Sup. Fig. 1a). This data is similar to IC_50_ values reported in TNBC [38, 42, 43], CRC [41], and glioblastoma [44] cells. IC_50_ values for hF31, hF37, hF44, and hT1 organoids were 20 nM, 75 nM, 28 nM, and 25 nM, respectively (Sup. Fig. 1b). Thus, TR-107 maintains low nanomolar potency across pancreatic cancer models.

**Figure 1.**
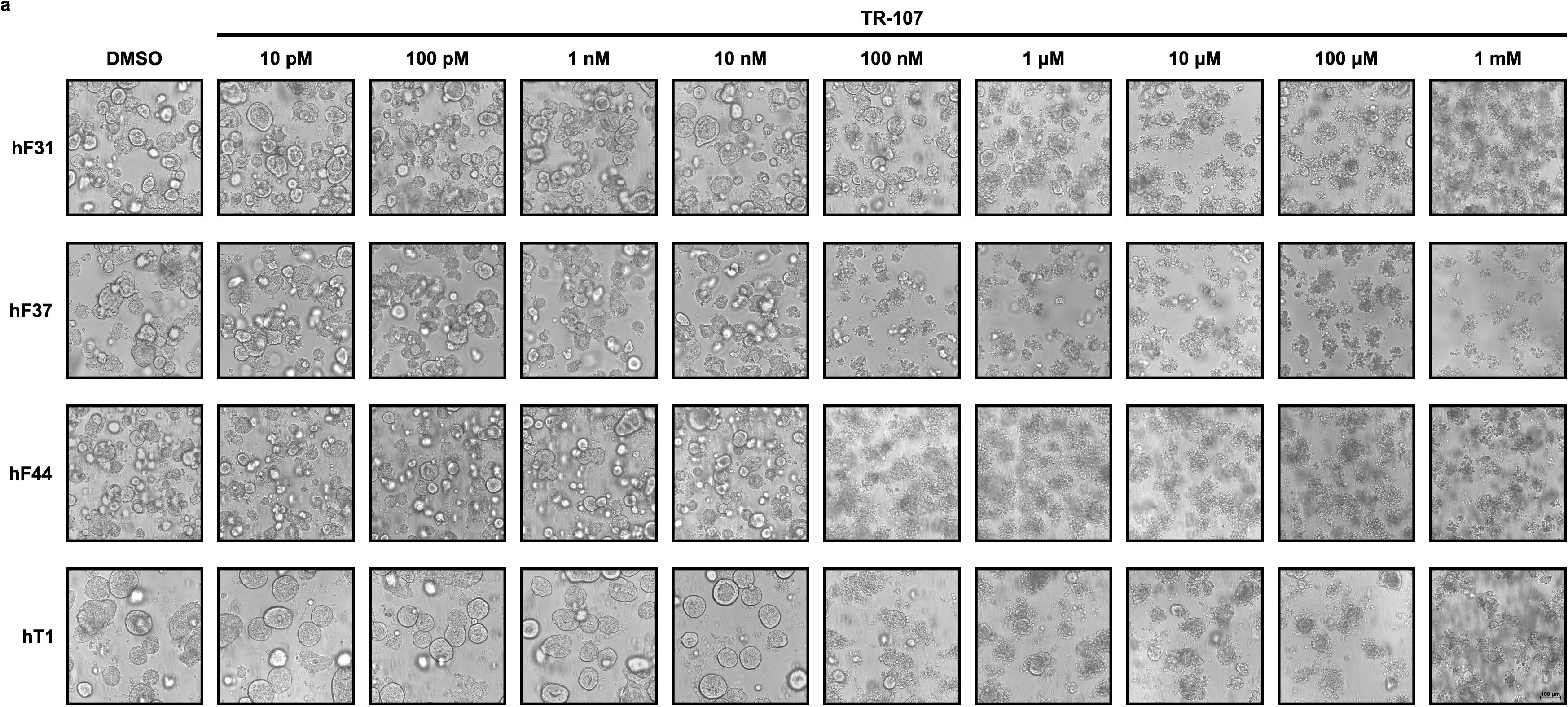
TR-107 exerts an inhibitory effect on growth in hF31, hF37, hF44, and hT1 PDAC organoids. **(A)** hF31, hF37, hF44, and hT1 PDAC organoids were seeded in Matrigel and treated with DMSO (vehicle) and with various concentrations of TR-107 for 72 hours. The figure displays representative images from a sample of n=5. The scale bar depicts 100 μm.

### TR-107 dysregulates essential mitochondrial proteins via ClpP proteolysis

Treatment with ONC201 and TR imipridone analogs, including TR-107, activate unregulated ClpP proteolysis in vitro [27]. This includes the dissociation and rapid degradation of ClpX in PDAC [22] and TNBC [42]. PDAC cell lines were treated with 10 nM, 50 nM, or 1 µM TR-107 or vehicle control (0.1% DMSO) for 24 hours, and immunoblot analysis of ClpX, and β-Actin was performed. A decrease in ClpX expression was observed in all four cell lines with the most significant effect between 10 nM and 50 nM TR-107 (Sup. Fig. 1c, 1d). With these results, organoids were seeded in Matrigel and treated with 50 nM TR-107 or vehicle control (0.1% DMSO) for 24 hours. Immunoblot confirmed downregulated ClpX expression by TR-107 in the four PDAC organoids (Fig. 2a, 2b). The mitochondrial transcription factor TFAM (mtFA) was also dysregulated by TR-107 in hF44, hF37, and hT1 (Fig. 2a, 2b) indicating aberrant mitochondrial function.

**Figure 2.**
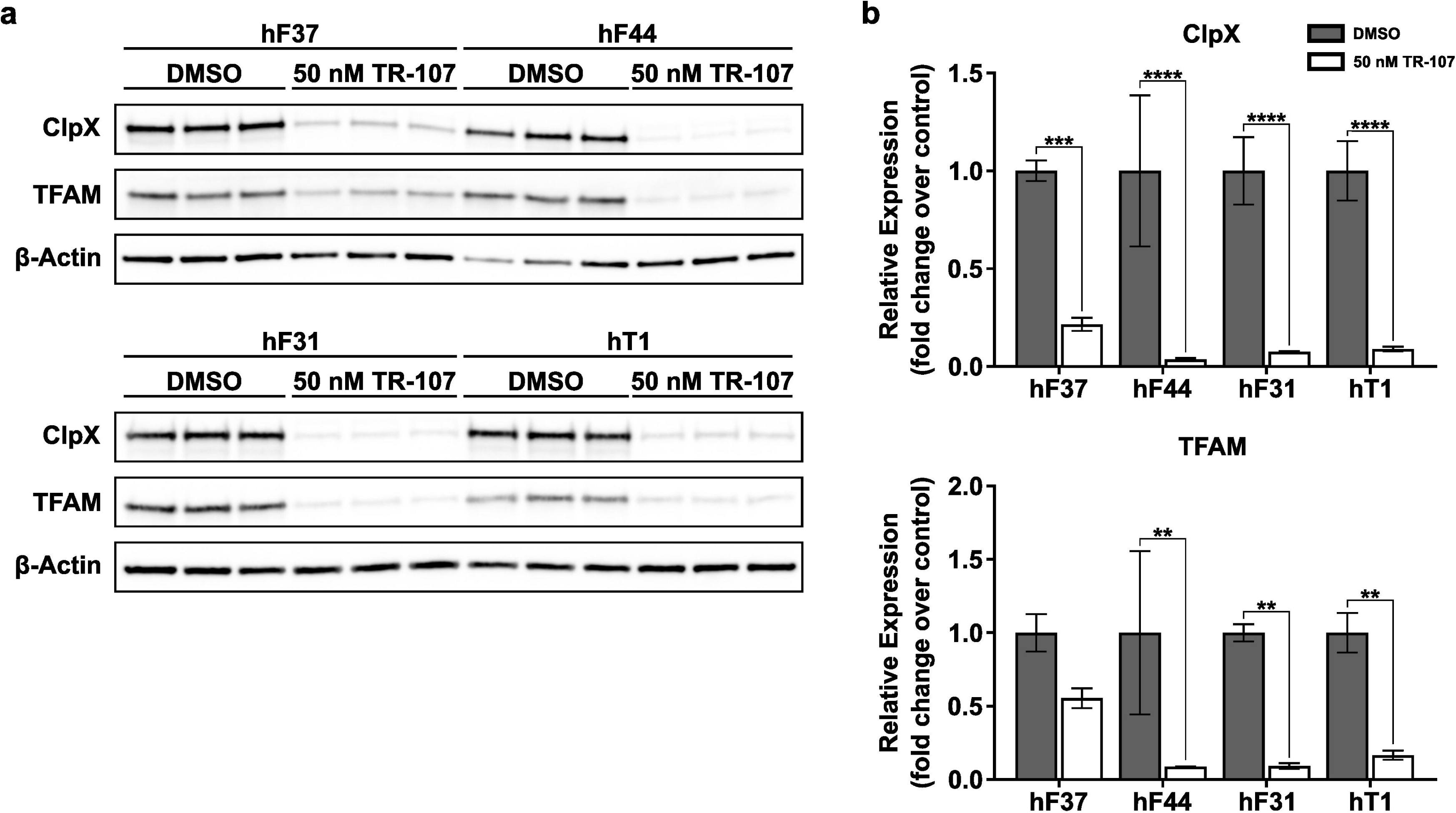
Mitochondrial ClpX and TFAM expression are inhibited a low-nanomolar concentration of TR-107. hF31, hF37, hF44, and hT1 PDAC organoids were seeded in Matrigel and treated with DMSO (vehicle) and 50 nM TR-107. Twenty-four hours after treatment, total protein was collected, and **(A)** immunoblot was performed for ClpX, TFAM, and β-Actin. **(B)** Densitometry was measured with ImageJ and Student’s t-test was conducted between treatments using GraphPad Prism with **p < 0.01, ***p < 0.001, ****p < 0.0001. Data are presented as mean ± SD (n=3).

### hF37 PDAC organoid transcriptome is altered by TR-107

To assess the effect of TR-107 on PDAC transcriptome, hF37 organoids were seeded in Matrigel and treated with 50 nM TR-107 or vehicle control (0.1% DMSO). Total RNA was extracted after 24- and 48-hours treatment, and bulk RNA-seq was performed. Analysis of both timepoints demonstrated little variance between 24 hours of TR-107 and the vehicle controls (Sup. Fig. 2a, 2b). Functional gene enrichment analysis further confirmed unsubstantial treatment response at this timepoint (Sup. Fig. 2c). Therefore, analysis of the 48-hour treatment was conducted separately. A difference between TR-107 and the respective vehicle control within the principal component analysis (Fig. 3a) and sample-to-sample heat map (Fig. 3b) implied a strong treatment effect on transcription at 48 hours. Furthermore, a high degree of concordance was observed within each cohort. Functional gene enrichment analysis of the top 40 dysregulated genes included mitochondrial ribosomal RNA (*MT-RNR1*, *MT-RNR2*), cellular and mitochondrial metabolism (*MTHFD2*, *SLC3A2*, *SLC7A5*, *SLC25A25*, *SLC7A1, SLC7A11*), and cellular stress (*GDF15*, *S100P*, *ATF4*) (Fig. 3c).

**Figure 3.**
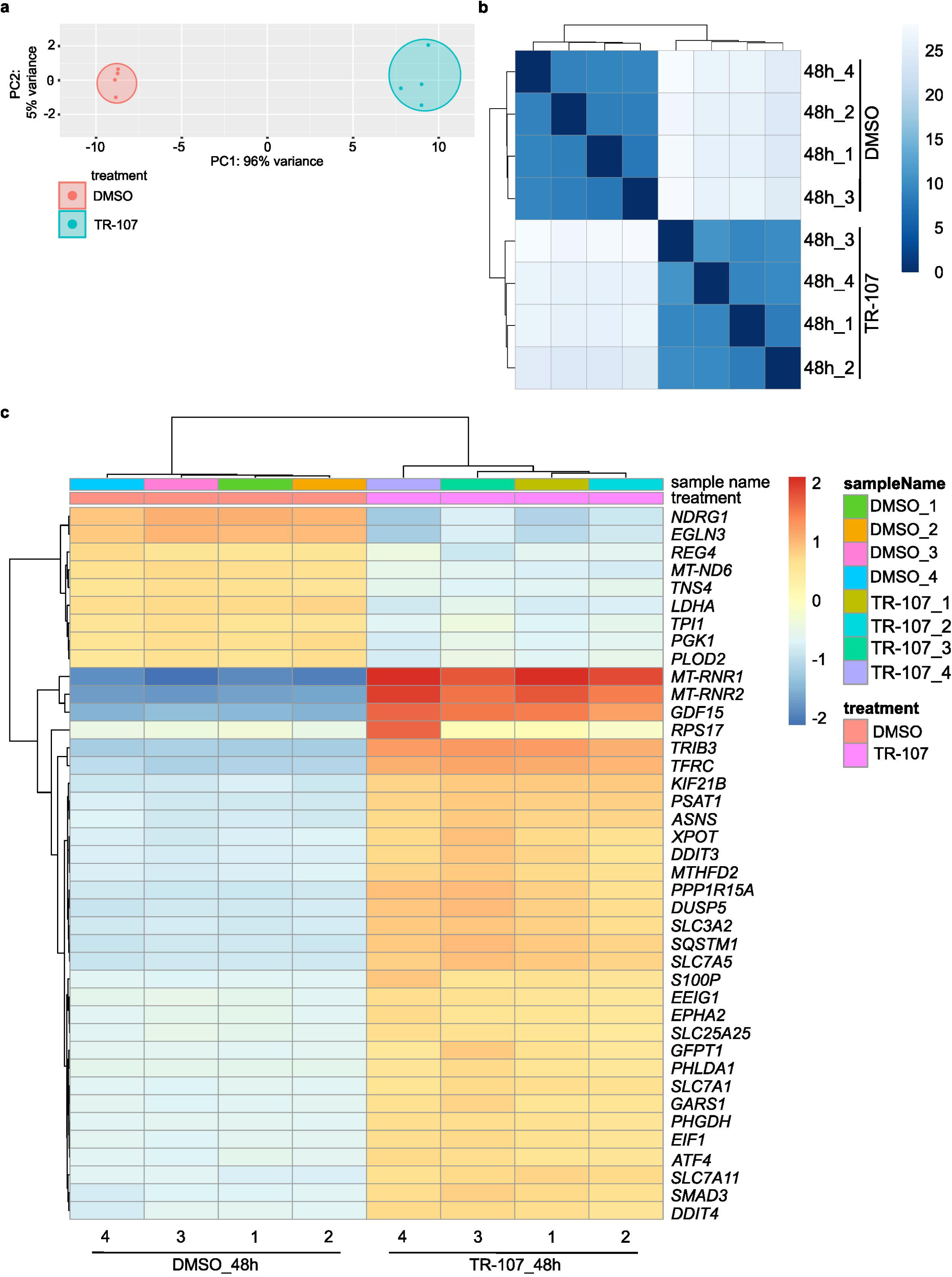
TR-107 treatment impacts the transcriptomic profile of PDAC organoids. hF37 PDAC organoid was seeded in Matrigel and treated with DMSO (vehicle) and with a 50 nM concentration of TR-107. Forty-eight hours after treatment, total RNA was extracted for bulk RNA-seq. **(A)** Principal component analysis (PCA) of hF37 PDAC organoid treated with DMSO and TR-107 for 48 hours. **(B)** Sample-to-sample heatmap of hF37 PDAC organoid treated with DMSO and TR-107 for 48 hours. **(C)** Functional enrichment of activated and suppressed genes and pathways upon TR-107 treatment of hF37 PDAC organoid.

### Ferroptosis and metabolism are dysregulated by TR-107

Following RNA-sequencing analysis, Kyoto Encyclopedia of Genes and Genomes (KEGG) pathway analysis was conducted to elucidate dysregulated cellular and molecular pathways. Cell growth and death pathways including ferroptosis (Fig. 4a, Sup. Fig. 3b) and apoptosis (Fig. 4a) were activated by TR-107 in hF37. Ferroptosis, an iron-dependent form of regulated cell death, was of particular interest given the gene ratio magnitude in hF37 and our recent findings in CRC [41]. This pathway is tightly regulated by Integrated Stress Response (ISR) which is induced by ONC201 and TR imipridone analogs via the master transcription factor Activating Transcription Factor 4 (ATF4) [37, 38, 40, 41]. Indeed, *ATF4* is significantly upregulated after TR-107 treatment in hF37 (Fig. 3c). RT-PCR confirmed upregulation of *ATF4* after 24- (Fig. 4b) and 48-hours (Fig. 4e) treatment of TR-107 in all four PDAC organoids.

**Figure 4.**
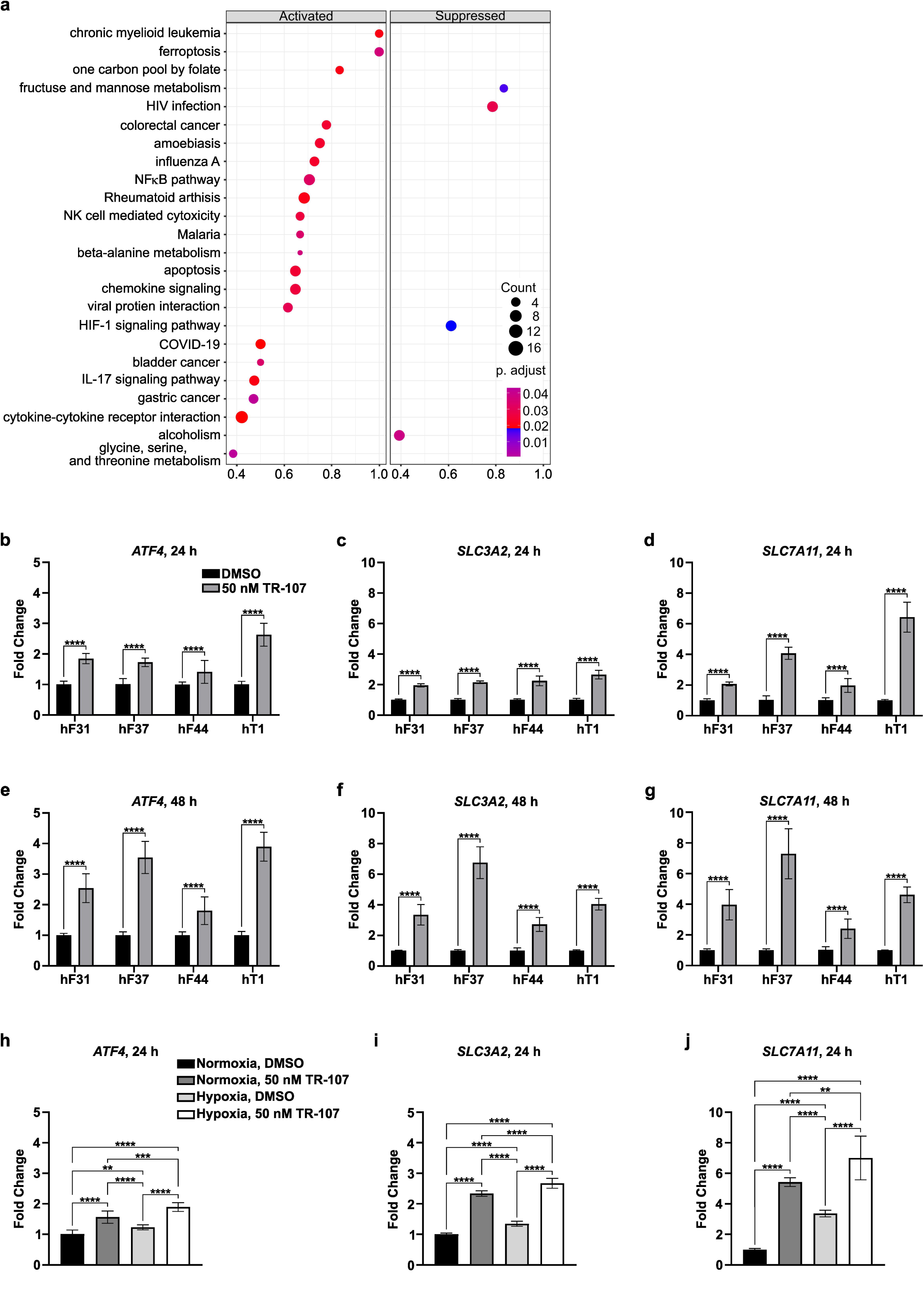
Treatment with TR-107 activates stress response and ferroptosis while reducing HIF1α signaling in hF37 PDAC organoids. hF31, hF37, hF44, and hT1 PDAC organoids were seeded in Matrigel and treated with DMSO (vehicle) and 50 nM TR-107, and RNA was collected 24- and 48-hour post-treatment. **(A)** Activated and suppressed pathways in hF37 PDAC organoid upon TR-107 treatment (48 hours). **(B-D)** RT-qPCR analysis of *ATF4*, *SLC3A2*, and *SLC7A11* in hF31, hF37, hF44, and hT1 PDAC organoids after 24 hours of TR-107 treatment, respectively. **(E-G)** RT-qPCR analysis of *ATF4*, *SLC3A2*, and *SLC7A11* in hF31, hF37, hF44, and hT1 PDAC organoids after 24 hours of TR-107 treatment, respectively. **(H-J)** RT-qPCR analysis of *ATF4*, *SLC3A2*, and *SLC7A11* in hF37 PDAC organoid under normoxia and hypoxia after 24 hours of TR-107 treatment, respectively. **(B-G)** Student’s t-test was conducted between treatments using GraphPad Prism. (H-J) 2-way ANOVA with multiple comparisons was conducted to compare treatments. Data are presented as mean ± SD values (n=3). **, p < 0.01; ***, p < 0.001; ****, p < 0.0001.

Amino acid metabolism pathways including beta-alanine; glycine, serine, and threonine (Fig. 4a); and glutathione synthesis (Sup. Fig. 3b) were also upregulated by TR-107. As previously described, metabolic reprogramming away from OXPHOS occurs because of the hypoxic and nutrient-poor microenvironment PDAC. This reliance on amino acid metabolism as a source of energy is exacerbated by TR-107-mediated mitochondrial dysfunction. The Solute Carrier (SLC) family of proteins facilitate diffusion, symport, and/or antiport of small molecules and ions across cell and organelle membranes. Solute Carrier Family 3 Member 2 (*SLC3A2*) and Solute Carrier Family 7 Member 11 (*SLC7A11*) are top activated transcripts in functional enrichment analysis (Fig. 3c) and ferroptosis pathway KEGG analysis (Sup. Fig. 3b). SLC3A2 and SLC7A11 heterodimerize to form the transmembrane System Xc− critical in the antiport exchange of extracellular L-cystine and intracellular L-glutamate. This results in glutathione synthesis required to protect against oxidative stress [45]. RT-PCR confirmed RNA-seq results for both *SLC3A2* and *SLC7A11* at 24- (Fig. 4c-d) and 48-hours (Fig. 4f-g) treatment of TR-107 in all four PDAC organoids. Upregulation of these Xc− antiporter subunits upon TR-107 treatment confirm a cellular response to increased oxidative stress caused by mitochondrial disfunction.

### TR-107 retains efficacy against PDAC cells in hypoxic conditions

A hallmark of PDAC is dense extracellular matrix secreted by activated PSCs resulting in a hypoxic, nutrient-poor TME. PDAC cells upregulate glycolysis and amino acid metabolism to maintain net energy production and promote survival in these conditions [19]. Transcriptomic analysis revealed the inhibition of HIF-1α signaling in hF37 organoids upon TR-107 treatment in normoxic conditions (Fig. 4a, Sup. Fig. 3a). To assess the efficacy of TR-107 in hypoxic conditions, organoids and cells were cultured in media containing CoCl_2_ (150 μM) for 12 hours followed by TR-107 treatment (50 nM) for 24, 48, and 72 hours while maintaining CoCl_2_-induced hypoxia. The normoxia treatment was performed as a control. Increased expression of *ATF4*, *SLC3A2*, and *SLC7A11* in hF37 treated with TR-107 in hypoxic culture was confirmed by RT-PCR (Fig. 4h-j). Thus, the effect of TR-107 on cellular stress response and amino acid metabolism is preserved in hypoxia.

Immunoblot analysis confirmed the induction of hypoxia by increased expression of HIF-1α in CoCl_2_-treated organoids (Fig. 5a-h) and cells (Sup. Fig. 4a-d) compared to the normoxia condition. HIF-1α levels were maintained across timepoints in cells treated with the DMSO vehicle control (Sup. Fig. 4a-d). However, HIF-1α expression was significantly reduced in organoids and cells treated with TR-107 (50 nM) compared to the vehicle control under hypoxic conditions. In PDAC cell lines, this was additionally observed across the three timepoints. ClpX and TFAM were downregulated in TR-107-treated organoids cultured under hypoxia compared to the vehicle control in hypoxia and normoxia conditions (Fig. 5a-h). Dysregulated ClpX expression was confirmed in the treated cell lines (Sup. Fig. 4a-d). Inhibition of ClpX and TFAM expression was comparable between normoxic and hypoxic conditions suggesting that TR-107-mediated proteolysis and aberrant mitochondrial function was not affected by hypoxia.

**Figure 5.**
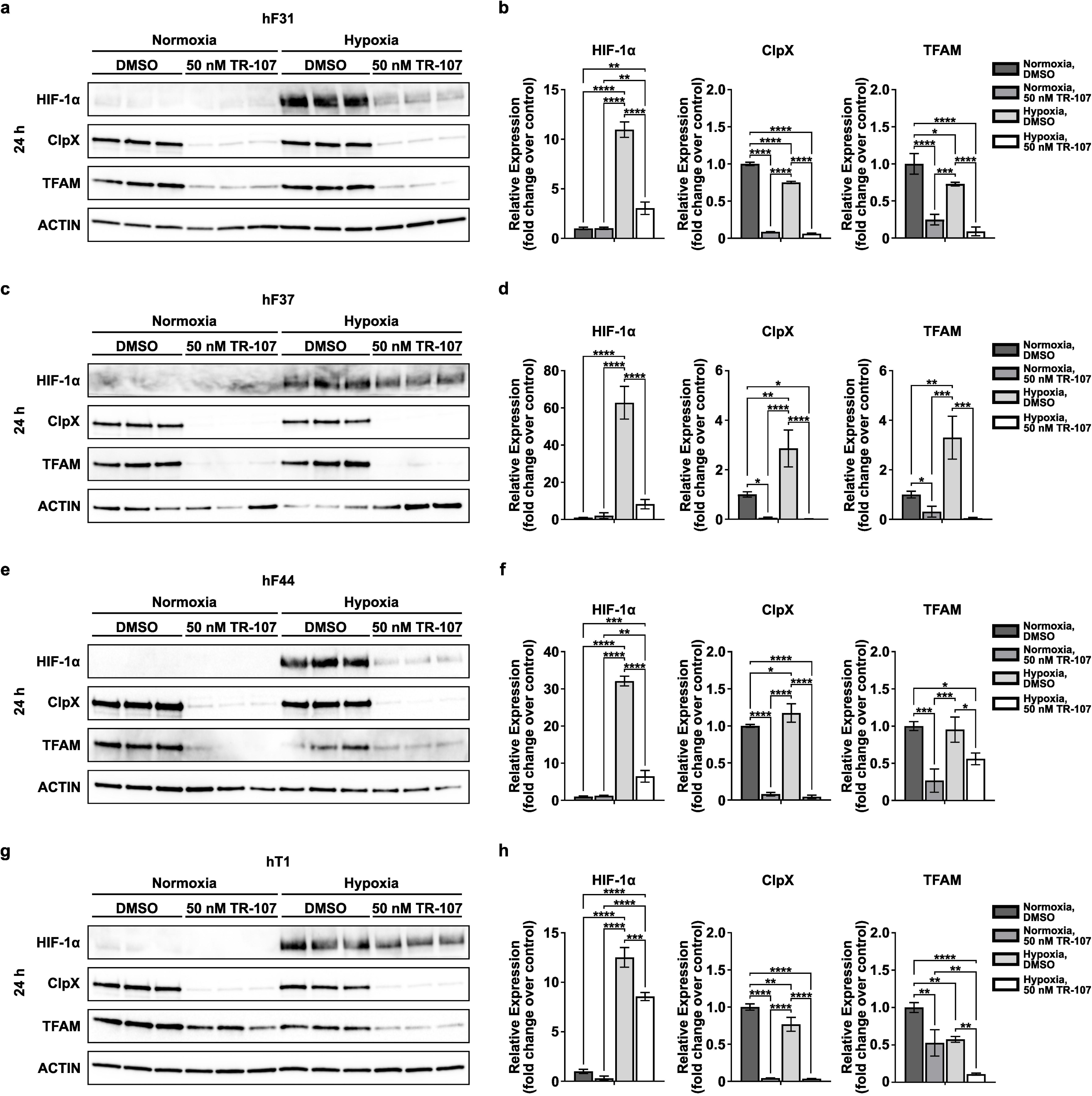
TR-107-mediated degradation of ClpX is preserved under hypoxic conditions in PDAC organoids. hF31, hF37, hF44, and hT1 PDAC organoids were seeded in Matrigel, and hypoxic conditions were induced for 24 hours, and treated with DMSO (vehicle) and with a 50 nM concentration of TR-107 for 24 hours, while maintaining CoCl_2_-induced hypoxia. **(A, C, E, and G)** immunoblot was performed for HIF1α, ClpX, TFAM, and β-Actin in hF31, hF37, hF44, and hT1 PDAC organoids, respectively. **(B, D, F, and H)** Densitometry was measured with ImageJ and ordinary One-way ANOVA was conducted between treatments using GraphPad Prism with * p < 0.05, **p < 0.01, ***p < 0.001, ****p < 0.0001. Data are presented as mean ± SD (n=3).

### TR-107 and Capivasertib synergize to dysregulate the AKT signaling pathway

To investigate the therapeutic sensitivity of PDAC organoids, the inhibitory efficacy of several compounds was evaluated including RMC-6236, Alpelisib, and Capivasertib. RMC-6236 (daraxonrasib) is a first-in-class RAS(ON) inhibitor targeting the active, GTP-bound form of RAS proteins (KRAS, HRAS, NRAS) across common mutations [46, 47]. It was recently approved for the treatment of adults with metastatic PDAC who previously failed systemic therapy [48]. While a decrease in viability was observed in the high nanomolar to low micromolar (100 nM – 10 µM) range for hF37 and hT1, a response was only seen at high micromolar concentration for hF31 and hF44 (Sup. Fig. 5) indicating low efficacy of RMC-6236 in some PDAC organoids. Calculation of the IC_50_ confirmed these viability results in the four organoids (Sup. Fig. 8a).

In PDAC, the PI3K/AKT/mTOR pathway is frequently hyperactivated by upstream KRAS signaling [49, 50]. This results in increased tumorigenesis, survival, and plasticity [51]. Thus, we next treated organoids with Alpelisib, a selective Phosphoinositide 3-kinase (PI3K) kinase inhibitor approved to treat hormone receptor (HR)+, HER2-advanced breast cancer with a *PIK3CA* gene mutation [52]. We observed an effect on viability of all four organoids when treated with a low micromolar (1 µM – 100 µM) concentration of Alperlisib (Sup. Fig. 6) and this was corroborated by the IC_50_ curve (Sup. Fig. 8b).

**Figure 6.**
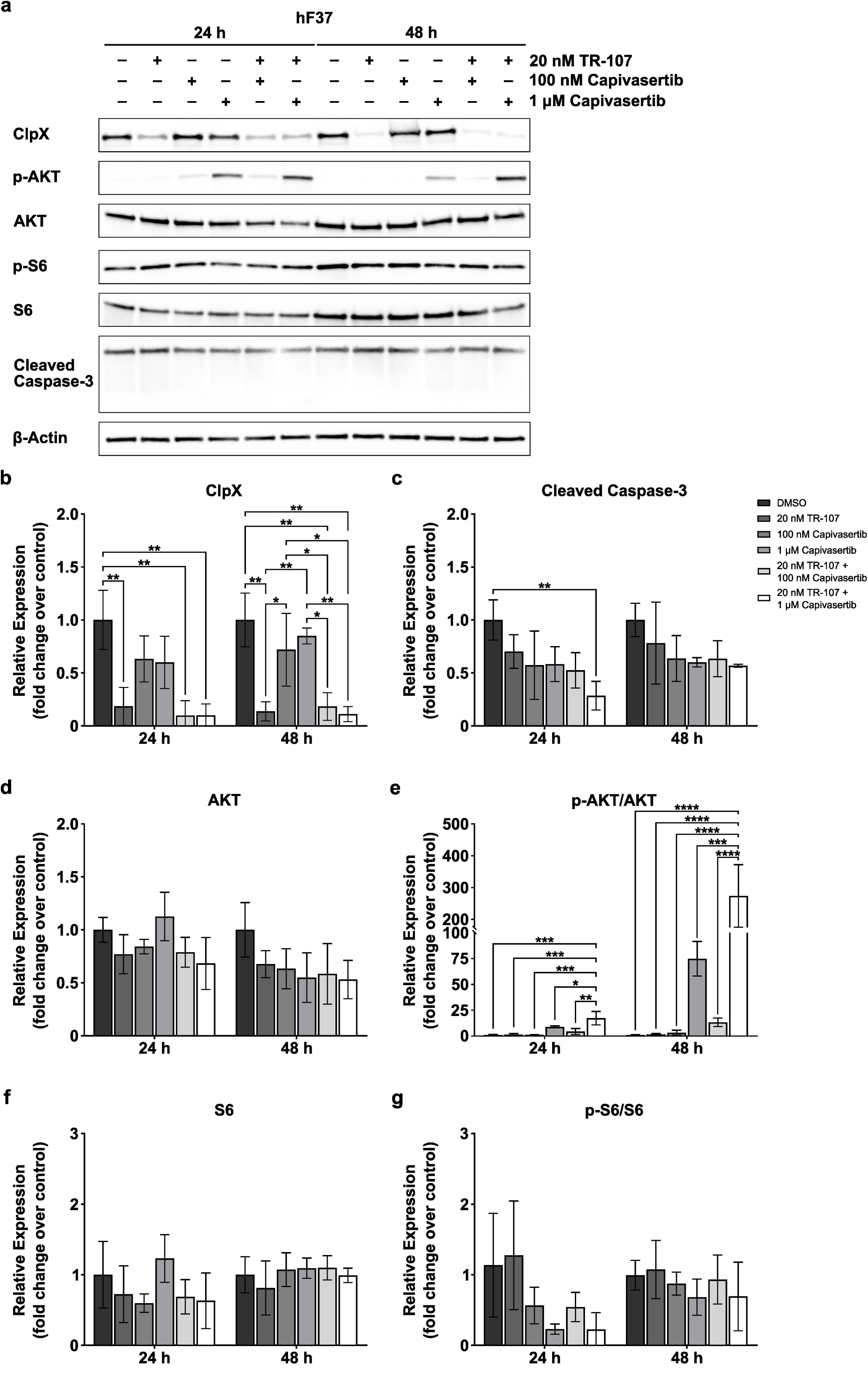
TR-107 and Capivasertib affected signaling pathway activity in the hF37 PDAC organoid. hF37 organoids were treated with DMSO (vehicle), 20 nM TR-107, 100 nM or 1 µM Capivasertib, and combinations thereof for 24 and 48 hours. **(A)** Immunoblots were performed for AKT, S6, phosphorylated counterparts, ClpX, Cleaved Caspase 3 and β-Actin as a loading control. **(B-G)** The densitometric analysis was conducted to quantify relative protein expression, normalized to β-actin control and non-phosphorylated proteins in samples after 24- and 48-hour post-treatment. Data are presented as mean ± SD values (n=3). *, *p* < 0.05; **, *p* < 0.01; ***, *p* < 0.001; ****, *p* < 0.0001.

Finally, we treated PDAC organoids with Capivasertib (brand name Truqap) currently approved for the treatment of HR+, HER2-advanced breast cancer with *PIK3CA*, *AKT1*, and/or *PTEN* mutations [53, 54]. Protein Kinase B (AKT) is downstream of PI3K and is upregulated by hyperactivated PI3K in approximately 60% of PDAC cases making it an alluring target for therapeutic inhibitors [55–57]. A decrease in viability was observed at 10 µM for hF37 and at 100 µM for hT1, but there was no apparent effect at the maximal concentration of 100 µM for hF31 and hF44 (Sup. Fig. 7). Furthermore, IC_50_ values could not be calculated for all organoids, either because treatment did not reach 50% inhibition or were ambiguous due to the nonlinear fit (Sup. Fig. 8c).

Given the spectrum of efficacy against inhibition of these compounds on the four PDAC organoids, synergy analysis was performed to determine if there was increased benefit of co-treatment with TR-107. hF37 and hF44 organoids were treated with TR-107 (10 nM or 20 nM or 30 nM or 40 nM) and RMC-6236 (5 nM or 10 nM or 20 nM), Alpelisib (50 nM or 100 nM or 1 µM), or Capivasertib (50 nM or 100 nM or 1 µM) for 72 hours. The greatest synergistic effect on inhibition was observed with co-treatment of TR-107 and Capivasertib (CAP) in hF37 (Sup. Fig. 9c) and hF44 (Sup. Fig. 10c). TR-107 and Alpelisib yielded a lesser degree of synergy (Sup. Fig. 9b, Sup. Fig. 10b) whereas TR-107 and RMC-6236 co-treatment resulted in additive or no additional effect on inhibition (Sup. Fig. 9a, Sup. Fig. 10a).

Immunoblot analysis was conducted to investigate the effect of synergistic co-treatment of TR-107 and Capivasertib on targeted downstream signaling pathways. hF37 and hF44 organoids were treated with DMSO (vehicle), 20 nM or 30 nM TR-107, 100 nM or 1 µM Capivasertib, and all combinations for 24 and 48 hours. ClpX was downregulated in all TR-107 treatment conditions in hF37 (Fig. 6a, b) and hF44 (Sup. Fig. 11a, b) at 24 and 48 hours. Singlet or co-treatment with Capivasertib had no effect on ClpX expression. Conversely, Capivasertib elicited a dose-dependent increase of phosphorylated-AKT in hF37 (Fig. 6a, e) and hF44 (Sup. Fig. 11a, e). Capivasertib does not inhibit AKT phosphorylation or total expression, but instead competitively binds to the ATP-binding pocket to block downstream signaling [58]. Therefore, cells paradoxically upregulate AKT phosphorylation in an attempt to maintain targeted effects [59, 60]. Singlet or co-treatment with TR-107 had no effect on p-AKT expression. While Cleaved Caspase-3 decreased with 30 nM TR-107 and 1 μM Capivasertib co-treatment at 24 hours in hF44 (Sup. Fig. 11a, c), this was not observed at 48 hours or in hF37 at either timepoint (Fig. 6a, c). No appreciable effect on S6 (total or phosphorylated) was seen in either organoid (Fig. 6a, f-g; Sup. Fig. 11a, f-g).

## Discussion

Presently, we establish the robust in vitro efficacy of the novel imipridone analog TR-107 in four PDAC organoid lines and four well-characterized cell lines. Inhibition of viability was reported at low-nanomolar concentrations (Fig. 1, Sup. Fig. 1a-b), and dose-dependent (10 nM, 50 nM, 1 μM) inhibition of ClpX was induced by TR-107 after 72 hours of treatment (Fig. 2, Sup. Fig. 1c-d). This potency against viability and ClpX expression in PDAC is consistent with that of other cancers [38, 41–44]. Unregulated mitochondrial ClpP protease activity after TR-107 treatment results from dissociation of ClpX from the ClpXP complex [38]. The resulting mitochondrial dysfunction contributes to aberrant cellular respiration, loss of homeostasis, and increased integrated stress response [41, 42].

A hallmark of PDAC is fibrosis driven by activated PSCs of the TME [19]. The resulting dense stroma creates an ischemic environment with minimal vasculature and a nutrient-deplete microenvironment. Metabolic reprogramming allows oncogenic cells to survive and proliferate under these conditions, while challenges to therapeutic delivery and chemoresistance persist [16–19]. We show that TR-107 treatment altered the hF37 transcriptome (Fig. 3c), specifically attenuating ferroptosis and amino acid metabolism while suppressing HIF-1α signaling (Fig. 4a). Mitochondrial stress and dysregulated OXPHOS caused by TR-107-induced ClpP mitochondrial proteolysis trigger ISR. Consequently, upregulated ATF4 transcriptionally activates components of redox and glutathione synthesis, including the cysteine/glutamate antiporter subunits SLC3A2 [61] and SLC7A11 [62–64]. Indeed, transcriptome analysis showed increased expression of *ATF4*, *SLC3A2*, and *SLC7A11* after TR-107 treatment (Fig. 3c), which was confirmed by RT-PCR (Fig. 4b-g). While pathway analysis indicates the involvement or perturbation of molecular pathways, it does not conclusively confirm that such pathways are activated or inhibited. We previously demonstrated that TR-107 induces ferroptosis in CRC [41]. However, ATF4 can paradoxically inhibit or promote ferroptosis in the context of ISR, redox imbalance, and dysregulated amino acid metabolism [65]. Future studies evaluating the effect of TR-107 on reactive oxygen species and antioxidant expression, including GPX4, should be conducted to characterize ATF4-mediated ferroptosis in PDAC.

To promote survival and proliferation in a hypoxic, nutrient-poor environment, PDAC cells upregulate glycolysis and amino acid metabolism. In normoxic culture, TR-107 inhibited the HIF-1α signaling pathway (Fig. 4a). This could render PDAC cells susceptible to nutrient and energy starvation and sensitize them to treatment. We then confirmed that TR-107 maintained efficacy against ClpX and mitochondrial function in a hypoxic environment (Fig. 5a-h, Sup. Fig. 4a-d). Consistent with transcriptomic analysis, TR-107 suppressed HIF-1α expression in hypoxia compared with the TR-107 and normoxia control. Recently, PDAC cells have been shown to induce activated PSCs under hypoxia. Inflammatory cytokines IL-1 and TNF-α derived from PDAC cells stabilized HIF-1α expression in CAF cells, consequently transforming them from a myofibroblastic phenotype to an inflammatory phenotype [66]. TR-107 may mitigate this PSC activation through HIF-1α inhibition and aberrant cytokine production. Hence, the effect of TR-107 on cytokine production in PDAC with/without PSC co-culture is of future interest.

Just as metabolic remodeling of cancer cells and cancer-associated fibroblasts confers chemoresistance, dysregulation of cellular metabolism may resensitize tumors to conventional therapies or novel compounds. After measuring the inhibitory effect of RMC-6236, Alpelisib, and Capivasertib alone, we conducted synergy studies in co-treatment with TR-107 in two PDAC organoids. Synergy was observed between TR-107 and Aleplisib and TR-107 and Capivasertib. We further studied TR-107 and Capivasertib due to Capivasertib’s function as an AKT inhibitor. As expected, the two compounds-maintained efficacy against their known targets (ClpX and AKT) when used as monotherapy or combinatorial treatment. Additionally, p-AKT increased in a dose-dependent manner when co-treated with TR-107. Thus, TR-107 has synergistic potential, and co-treatment may sensitize PDAC to other therapeutics.

## Methods

### Cell lines and compounds

Human PDAC cell lines PANC-1 (ATCC, CRL-1469, RRID:CVCL_0480) and MIA PaCa-2 (ATCC, CRL-1420, RRID:CVCL_0428) were cultured in DMEM medium (Thermo Fisher Scientific, 10-013-CV); BxPC-3 (ATCC, CRL-1687, RRID:CVCL_0186) was cultured in RPMI-1640 medium (Thermo Fisher Scientific, 10-040-CV); and Capan-2 (ATCC, HTB-80, RRID:CVCL_0026) was cultured in McCoy’s 5a medium (Thermo Fisher Scientific, 10-050-CV). All media were supplemented with fetal bovine serum (10% v/v; Peak Serum, PS-FB3) and penicillin/streptomycin (1% v/v; Gibco, 15140-122). Human pancreatic cancer organoids (hF31, hF37, hF33, and hT1) were obtained from Stony Brook Cancer Center Biobank and Cold Spring Harbor Laboratory Biobank and grown according to the previous publications [67, 68]. All organoids and cell lines were maintained at 37°C and CO_2_ (5%), and all experiments were conducted at a cell passage of less than 30.

To replicate hypoxic conditions and induce HIF-1α expression, cells were cultured in media containing CoCl_2_ (150 μM) for 12 hours [69]. CoCl_2_ stabilizes HIF-1α, thus preventing degradation under normoxic conditions [70–72]. Hypoxia was maintained during successive treatment with TR-107 (50 nM). A normoxic (media-only) control was included in all experiments.TR-107 compound was synthesized and provided by Madera Therapeutics, LLC. RMC-6236 (E1597), Alpelisib (NC0421898), and Capivasertib (501947955) were purchased from Selleckchem.com. All compounds were reconstituted in dimethyl sulfoxide (DMSO; Fisher Bioreagents, BP231-100). A vehicle control of DMSO (0.1% v/v) was used in all experiments.

### Cell viability assay

BxPC-3, Capan-2, MIA PaCa-2, and PANC-1 cells were seeded at 1.0 x 10^3^ cells/well in opaque flat-bottom 96-well plates (Corning, 3917) with 100 µl of respective media/well. Twenty-four hours post-seeding, cells were treated with vehicle control (0.1% DMSO) or log-fold concentrations of TR-107 for 72 hours at 37°C and CO_2_ (5%). hF31, hF37, hF44, and hT1 organoids were seeded in Matrigel (50% Matrigel:50%dPBS) in a 96-well microplate (Corning) in 100 μL of appropriate media and incubated for 24 hours. Following incubation, the growth media were supplemented with 100 μL vehicle control (0.1% DMSO) or various concentrations of TR-107, or RMC-6236, or Alpelisib, or Capivasertib for 72 hours at 37°C and CO_2_ (5%). Cell viability was assessed using the Cell Titer-Glo luciferase assay (Promega, G7571), and luminescence was measured using a SpectraMax M3 microplate reader (Molecular Devices). Half-maximal inhibitory concentrations (IC_50_) values were calculated using log(inhibitor) vs response – Variable slope (four parameters) nonlinear fit in GraphPad Prism (GraphPad Software version 10.4.1).

### Live organoids imaging

hF31, hF37, hF44, and hT1 PDAC organoids were seeded in the Matrigel (50% Matrigel:50%dPBS) in a 96-well microplate (Corning) in 100 μL of appropriate media and incubated for 24 hours. Following incubation, the growth media were supplemented with 100 μL vehicle control (0.1% DMSO) or log-fold concentrations of TR-107 or RMC-6236 or Alpelisib, or Capivasertib for 72 hours at 37°C and CO_2_ (5%). Images of the organoids were taken 24-, 48-, and 72-hour post-treatment using Nikon ECLIPSE Ti2 inverted microscope. The images were processed using NIS-Elements Advanced Research 5.21.03 software (RRID:SCR_027181).

### Western Blot analysis

BxPC-3, Capan-2, MIA PaCa-2, and PANC-1 cells were seeded at 7.5 x 10⁴ cells/well in a 6-well plate format (2.0 mL respective media/well). Twenty-four hours post-seeding, cells were treated with vehicle control (0.1% DMSO) or concentrations of TR-107 (10 nM, 50 nM, or 1 µM) and incubated at 37°C and CO_2_ (5%).

hF31, hF37, hF44, and hT1 PDAC organoids were seeded Matrigel in a 6-well plate (Corning) in 2.5 mL of appropriate media and incubated for 24-48 hours as previously described [73]. Following incubation, the growth media were supplemented with 0.2% DMSO or various concentrations of TR-107 and Capivasertib. Organoids were washed with ice-cold DBPS and lysed in Laemmli buffer 24- and 48-hours post-treatment.

Cell lysates were heat-denatured total protein extracts and mass-separated by sodium dodecyl sulfate – polyacrylamide gel electrophoresis (SDS-PAGE), transferred to a nitrocellulose membrane, and blocked in non-fat milk (5% w/v in Tris-buffered saline containing 0.01% Tween 20 (TTBS)) for one hour at room temperature. Membranes were then incubated overnight at 4°C with primary antibodies against human ClpX (1:1000; Abcam, ab168338), TFAM (1:1000, Cell Signaling, cs-8076), phosphorylated AKT (1:1000, Thermo Fisher Scientific, 44-621G), AKT (1:000, Cell Signaling, cs-9272), phosphorylated S-6 (1:1000, Cell Signaling, cs-4856), S6 (1:1000, Cell Signaling, cs-2317), Cleaved Caspase 3 (1:1000, Cell Signaling, cs-9661), HIF-1α (1:1000, Cell Signaling, cs-36169), and human β-Actin (1:2000; Millipore Sigma, A1978). The next day, membranes were washed in TTBS and incubated with HRP-conjugated secondary antibodies, goat anti-mouse (Sigma-Aldrich, AP200P) or goat anti-rabbit (Jackson ImmunoResearch, 111-035-144) for one hour at room temperature. Chemiluminescence was detected by SuperSignal^TM^ West Pico PLUS Chemiluminescent Substrate (ThermoFisher Scientific, 34578) and imaged using Azure c400 Visible Fluorescent Imager (Azure Biosystems, 10147-218, RRID:SCR_028261). Relative band density was measured using ImageJ (version 1.8.0_345, National Institutes of Health) [74], normalized to respective β-Actin, and analyzed in GraphPad Prism (GraphPad Software version 10.0.2, RRID:SCR_002798).

### Bulk RNA-sequencing (RNA-seq)

Total RNA was extracted using the QIAGEN RNeasy Mini Kit (QIAGEN, 74104). RNA integrity number (RIN) was measured using High Sensitivity RNA TapeStation Assay and all samples had a RIN> 9. The cDNA library construction and paired-end 150 bp sequencing were performed on Illumina NovaSeq platforms by Novogene (Davis Lab, Sacramento, CA). Pair-end reads were aligned to the human genome (hg38) using HISAT2 (RRID: SCR_015530), and transcript counts were quantified with featureCounts (RRID: SCR_012919) [75]. Gene expression fold-change ≥ 2 and false discovery rate (FDR) < 0.05 were set as the threshold values for subsequent analysis. Downstream analysis was performed in R/RStudio [76] using DESeq (RRID: SCR_000154) [77], apelgm [78], ComplexHeatmap [79], EnhancedVolcano [80], and following the previously described workflow [81].

### Quantitative Reverse Transcription PCR (RT-qPCR

hF31, hF37, hF44, and hT1 PDAC organoids were seeded as described above and treated with vehicle control (0.1% DMSO) or TR-107 50 nmol/L. After 24 and 48 hours of treatment, RNA was isolated using the QIAGEN RNeasy Mini Kit (QIAGEN, 74104), and cDNA was synthesized using SuperScript IV VILO Master Mix (Thermo Fisher Scientific, 11756050). *ACTB* was used as the housekeeping gene. TaqMan primers used were as follows: *ATF4* (ID: Hs00909569_g1, 4453320), *SLC3A2* (ID: Hs00374243_m1, 4453320), and *SLC7A11* (ID: Hs00921938_m1, 4453320). TaqMan RTq-PCR quantification was performed using the QuantStudio 3 Real-Time PCR System (Applied Biosystems, A28567). Samples were run in technical triplicate in a 96-well plate format. Data were analyzed using the Comparative CT (ΔΔCT) method [82].

### Synergy analysis

Synergy assays were conducted similarly to the viability assay. Each point (mean ± standard error of the mean) represents the growth of compound-treated cells compared to vehicle-only-treated cells. The expected growth inhibition was calculated using the HSA independence model and overlaid on the experimental growth curve in GraphPad Prism for Windows version 10.4.1. The synergy analysis was performed using SynergyFinder 3.0 in R [83, 84]. The data are presented as a dose-response matrix and Bar Plot for Multi-drug Combination Dose-Response/Synergy scores.

### Statistical analysis

Statistical analysis was performed by One-Way ANOVA, Two-Way ANOVA with Tukey test for multiple comparisons, and Student’s t-test for two comparison groups. All statistical analyses were performed using GraphPad Prism (Version 10.0.2).

## Acknowledgements

The authors wish to acknowledge the Stony Brook University Genomics Core for assistance with RNA-seq. We thank Michael Giarrizzo for help with data analysis. This research was funded by the National Institutes of Health grant R01GM138520 to L.M.G., R01DK124342 to A.B.B., and the Stony Brook Cancer Center grant to A.B.B.

## Author Contributions

Conception: ABB, LMG, and EJI; design: ABB, LMG, EJI; acquisition: JFL, AM, CV, SK, MK; analysis: JFL, AM, CV, SK, MK, and ABB; interpretation of data: JFL, AM, CV, SK, MK, and ABB, LMG, and EJI; writing – original draft, review, and editing: JFL, AM, CV, SK, MK, and ABB, LMG, and EJI.

## Data Availability

RNA-seq data generated for this study have been deposited in NCBI’s Gene Expression Omnibus (GEO) (RRID: SCR_005012) and are accessible through the GEO Series accession number: GSE338954. All other data are available upon request from the corresponding author.

## Additional Information

Competing Interests Statement: EJI has an ownership interest in Madera Therapeutics, LLC. All other authors declare no conflicts of interest.

---

**Supplementary Figure 1. TR-107 inhibits PDAC cells and organoids viability in vitro at low nanomolar concentrations. (A)** BxPC-3, Capan-2, MIA PaCa-2, and PANC-1 PDAC cells, and **(B)** hF31, hF37, hF44, and hT1 PDAC organoids were treated with log-fold serial dilutions of and TR-107. CellTiter-Glo luminescent cell viability assay was conducted 72 hours post-treatment. Half-maximal inhibitory concentrations (IC_50_) were determined using log(inhibitor) vs response – Variable slope (four parameters) nonlinear fit in GraphPad Prism (GraphPad Software version 10.0.2). Data are presented as mean ± SD (n=6). **(C)** BxPC-3, Capan-2, MIA PaCa-2 and PANC-1 PDAC cells were treated with 0.1% DMSO (control) or 10 nM or 50 nM or 1 µM TR-107. Twenty-four hours after treatment, total protein was collected and **(C)** immunoblot was performed for ClpX and β-Actin. **(D)** Densitometry was measured with ImageJ and Student’s t-test was conducted between treatments using GraphPad Prism with **p < 0.01, ***p < 0.001, ****p < 0.0001. Data are presented as mean ± SD (n=3).

**Supplementary Figure 2. TR-107 treatment impacts the transcriptomic profile of PDAC organoids.** hF37 PDAC organoid was seeded in Matrigel and treated with DMSO (vehicle) and with a 50 nM concentration of TR-107. Twenty-four and forty-eight hours after treatment, total RNA was extracted for bulk RNA-seq. **(A)** Principal component analysis (PCA) of hF37 PDAC organoid treated with DMSO and TR-107 for 24 and 48 hours. **(B)** Sample-to-sample heatmap of hF37 PDAC organoid treated with DMSO and TR-107 for 24 and 48 hours. **(C)** Functional enrichment of activated and suppressed genes and pathways upon TR-107 treatment of hF37 PDAC organoid for 24 and 48 hours.

**Supplementary Figure 3. Examples of pathways affected by TR-107 treatment in hF37 PDAC organoid.** HIF1 signaling pathway **(A)** and ferroptosis pathway **(B)** with downregulated and upregulated genes upon 48 hours of treatment with TR-107 as identified by KEGG analysis.

**Supplementary Figure 4. TR-107-mediated degradation of ClpX is preserved under hypoxic conditions in PDAC cell lines**. BxPC-3, Capan-2, MIA PaCa-2, and PANC-1 PDAC cell lines were seeded in a 6-well plate format, and hypoxia conditions were induced for 24 hours, followed by treatment with DMSO (vehicle) and 50 nM concentration of TR-107 for 24 hours, while maintaining CoCl_2_-induced hypoxia. Immunoblot and densitometry were performed for HIF1α, ClpX, and β-Actin in **(A)** BxPC-3, **(B)** Capan-2, **(C)** MIA PaCa-2, and **(D)** PANC-1 PDAC cell lines, respectively. Densitometry was measured with ImageJ and ordinary One-way ANOVA was conducted between treatments using GraphPad Prism with * p < 0.05, **p < 0.01, ***p < 0.001, ****p < 0.0001. Data are presented as mean ± SD (n=3).

**Supplementary Figure 5. RMC-6236 exerts an inhibitory effect on growth in hF31, hF37, hF44, and hT1 PDAC organoids. (A)** hF31, hF37, hF44, and hT1 PDAC organoids were seeded in Matrigel and treated with DMSO (vehicle) and with various concentrations of TR-107 for 72 hours. The figure displays representative images from a sample of n=5. The scale bar depicts 100 μm.

**Supplementary Figure 6. Alpelisib exerts an inhibitory effect on growth in hF31, hF37, hF44, and hT1 PDAC organoids. (A)** hF31, hF37, hF44, and hT1 PDAC organoids were seeded in Matrigel and treated with DMSO (vehicle) and with various concentrations of TR-107 for 72 hours. The figure displays representative images from a sample of n=5. The scale bar depicts 100 μm.

**Supplementary Figure 7. Capivasertib exerts an inhibitory effect on growth in hF31, hF37, hF44, and hT1 PDAC organoids. (A)** hF31, hF37, hF44, and hT1 PDAC organoids were seeded in Matrigel and treated with DMSO (vehicle) and with various concentrations of TR-107 for 72 hours. The figure displays representative images from a sample of n=5. The scale bar depicts 100 μm.

**Supplementary Figure 8. RMC-6236, Alpelisib, and Capivasertib inhibit the viability of multiple PDAC organoids.** hF31, hF37, hF44, and hT1 PDAC organoids were treated with variable concentrations of RMC-6236 **(A)**, Alpelisib **(B)**, and Capivasertib **(C)** for 72 hours, and cell viability was measured using 3D Cell Titer-Glo. The results are shown as mean ±SD (n=5) and normalized to the control (DMSO). Half-maximal inhibitory concentrations (IC_50_) were determined using log(inhibitor) vs response – Variable slope (four parameters) nonlinear fit in GraphPad Prism (GraphPad Software version 10.0.2).

**Supplementary Figure 9. Synergy analysis between TR-107 and RMC-6236 or Alpelisib, or Capivasertib in hF37 PDAC organoid.** hF37 PDAC organoids were treated with TR-107 10 nM or 20 nM or 30 nM or 40 nM and **(A)** RMC-6236 of 5 nM or 10 nM or 20 nM, **(B)** Alpelisib of 50 nM or 100 nM or 1 µM, **(C)** Capivasertib of 50 nM or 100 nM or 1 µM for 72 hours. Synergy analysis was performed using SynergyFinder using ZIP, Loewe, HSA, and Bliss algorithms. Data are presented as a dose-response matrix and Bar Plot for Multi-drug Combination Dose-Response/Synergy scores.

**Supplementary Figure 10. Synergy analysis between TR-107 and RMC-6236 or Alpelisib, or Capivasertib in hF44 PDAC organoid.** hF37 PDAC organoids were treated with TR-107 10 nM or 20 nM or 30 nM or 40 nM and **(A)** RMC-6236 of 5 nM or 10 nM or 20 nM, **(B)** Alpelisib of 50 nM or 100 nM or 1 µM, **(C)** Capivasertib of 50 nM or 100 nM or 1 µM for 72 hours. Synergy analysis was performed using SynergyFinder using ZIP, Loewe, HSA, and Bliss algorithms. Data are presented as a dose-response matrix and Bar Plot for Multi-drug Combination Dose-Response/Synergy scores.

**Supplementary Figure 11. TR-107 and Capivasertib affected signaling pathway activity in the hF44 PDAC organoid.** hF44 organoids were treated with DMSO (vehicle), 30 nM TR-107, 100 nM or 1 µM Capivasertib, and combinations thereof for 24 and 48 hours. **(A)** Immunoblots were performed for AKT, S6, phosphorylated counterparts, ClpX, Cleaved Caspase 3 and β-Actin as a loading control. **(B-G)** The densitometric analysis was conducted to quantify relative protein expression, normalized to β-actin control and non-phosphorylated proteins in samples after 24- and 48-hour post-treatment. Data are presented as mean ± SD values (n=3). *, *p* < 0.05; **, *p* < 0.01; ***, *p* < 0.001; ****, *p* < 0.0001.

